# Evaluation of a Functional Portable Hemodialysis Machine, XKIDNEY

**DOI:** 10.1101/2024.10.25.620359

**Authors:** Jake K. Lee, Dilara Derya, Hanui Yang, Patrick C. Lee

**Affiliations:** Exorenal Inc.; University of Toronto

**Keywords:** Portable Hemodialysis, Hemodialysis, Renal Care, End-Stage Renal Disease, Chronic Kidney Disease

## Abstract

Chronic kidney failure and end-stage renal disease (ESRD) are increasing globally, but home hemodialysis offers a critical means of improving patient quality of life and reducing national healthcare costs. This study investigates the performance of a portable hemodialysis (HD) device named XKIDNEY. The efficacy and safety of the device were determined with focus on solute clearance, fluid balancing accuracy, and blood cell integrity in an ex-vivo test environment. A comparison of the performance of XKIDNEY and a conventional HD unit showed that XKIDNEY had superior balancing accuracy without any technical malfunction. In addition, low molecular weight solutes, such as urea and creatinine, were effectively removed. These findings demonstrate that XKIDNEY enhances treatment portability without compromising performance and indicate XKIDNEY is a feasible home hemodialysis option.

---

Kidney failure is a rapidly spreading global disease that is increasing at an annual rate of ∼9.5%.^1,2^ Currently, in-center HD, which requires patients to attend 4-hour hospital sessions three times per week, accounts for 89% of all kidney failure therapies.^3^

Lack of patient autonomy over treatment scheduling poses significant burdens on patients and their families, such as extended commuting times, lengthy treatment durations, mobility, social interaction limitations due to rigid dialysis schedules, and strict dietary restrictions due to insufficient dialysis dose, which collectively degrade quality of life.^4^ Furthermore, nearly 50% of HD patients experience cardiovascular complications,^5,6^ and the five-year survival rate of these patients remains at only 50%.^7,8^

Dialysis becomes a lifelong necessity when they don’t have a kidney transplant. The high cost of in-center HD has emerged as a critical societal issue largely due to the financial burden it places on national healthcare systems responsible for covering the dialysis costs.^9^ In the United States, dialysis patients comprise only 1% of the total Medicare population but consume 7% of total Medicare expenditure.^8^ In South Korea, a policy implemented in response to the rapidly increasing financial demands of dialysis treatment has frozen the cost of HD for 20 years.^10,11^ These examples underscore the broader economic strain imposed by long-term dialysis care on healthcare systems.

The challenges associated with traditional dialysis could be substantially alleviated by enabling patients to perform dialysis at home, as was underscored by a U.S. government intention to increase the home dialysis rate.^9,12^ Currently, peritoneal dialysis is usually used for home dialysis. However, the number of patients requiring a transition to HD is rising due to complications such as peritonitis and reductions in peritoneal function.^12-14^ Thus, an increase in home HD appears inevitable. In view of the potential benefits of home HD, in terms of enhancing quality of life and reducing national healthcare expenditure, this shift in treatment methodology represents a progression in dialysis care.^15,16^

Overcoming the limitations of existing technologies and providing user-friendly home HD equipment would revolutionize end-stage renal disease (ESRD) care delivery, increase patient independence, and reduce the strain on healthcare systems. The development of XKIDNEY addresses the challenges patients face and supports the activities of hospitals and national health insurance systems.^17,18^ XKIDNEY facilitates the transition of hospital-based dialysis to homes and considerably reduces the time and location-associated limitations traditionally associated with HD. XKIDNEY allows patients to undergo HD at any time in any place, and thus, greatly enhances treatment flexibility and accessibility.

In the present study, we investigated the hemodialytic efficiency of the XKIDNEY unit, including solute clearances and balancing accuracy, in an ex-vivo setting. The results obtained were compared with those of conventional HD unit, with an emphasis on molecular reductions, albumin loss, and blood cell damage. The feasibility of the portable XKIDNEY unit was confirmed by monitoring blood and dialysate flow stabilities during dialysis sessions.

## MATERIALS AND METHODS

The XKIDNEY HD system shown in FIG. 1 is designed for home and in-center use and markedly simplifies dialysis procedures. By adopting a novel, proprietary pumping technology, the essential functions of dialysate and e?uent pumps, the balancing chamber, and the ultrafiltration (UF) pump, all previously considered indispensable for traditional HD devices, were integrated into a single unit, which resulted in a remarkable reduction in the unit size and weight (a volume of ∼57 Liters and a weight of ∼30 kg). The unit enabled dialysate flow rates upstream and downstream of the hemodialyzer, which reflects fluid management, to be regulated more accurately.

**FIG. 1.**
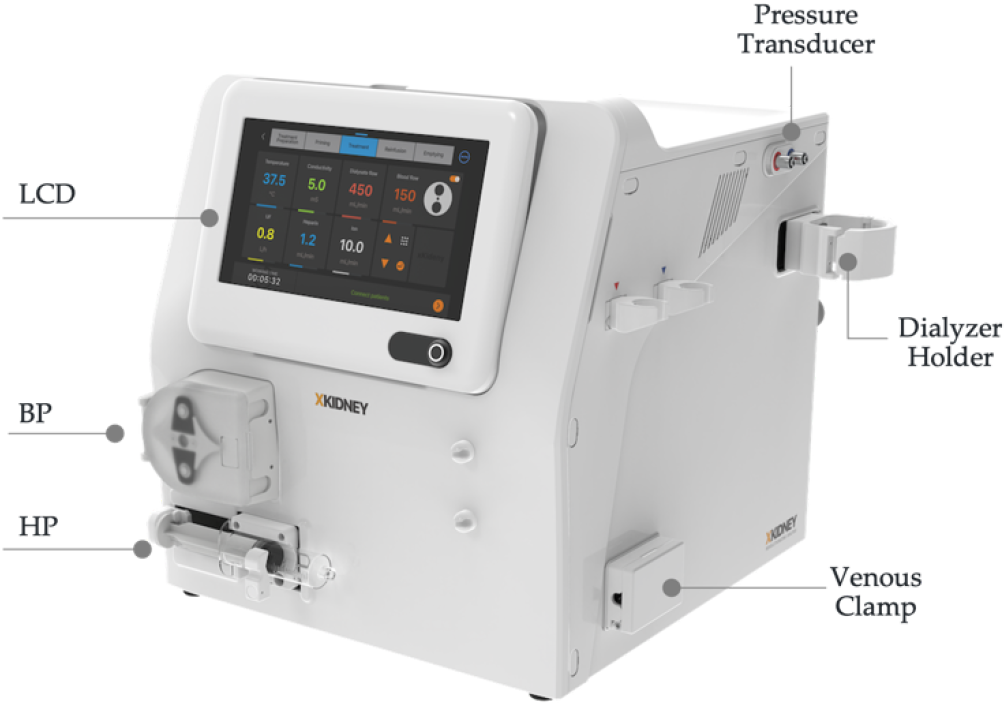
XKIDNEY (BP, blood pump; HP, heparin pump)

XKIDNEY is compatible with the various water sources commonly used for dialysis treatments, such as hospital-installed water treatment facilities or stand-alone portable reverse osmosis (RO) units, which provide unlimited amounts of ultrapure dialysate.

The device also provides greater treatment flexibility and dialytic performance. Specifically, XKIDNEY provides conventional high-flux HD using traditional acid- and bicarbonate-based dialysate, dialysate-free hemofiltration, and push/pull hemodiafiltration (HDF). The unit also utilizes standard consumables, such as normal blood tubing, concentrates, and a dialyzer.

Furthermore, the graphic touchscreen interface offers intuitive treatment instructions while monitoring patient progression. Thus, XKIDNEY improves ease of use and procedural convenience and safety. In addition, XKIDNEY features a more straightforward disinfection process and reduces manufacturing costs, which lower the cost of dialysis and provide cost savings for dialysis clinics and government healthcare systems.

### Hemodialytic Efficiency

Hemodialytic efficiency was investigated using two experimental groups – XKIDNEY and conventional dialysis groups. A schematic of the experimental circuit is provided in FIG. 2. An XKIDNEY unit was used for the XKIDNEY group, and an NCU-18 (Nipro, Japan) unit was used for the conventional dialysis group. Other experimental conditions were identical for the two groups.

**FIG. 2.**
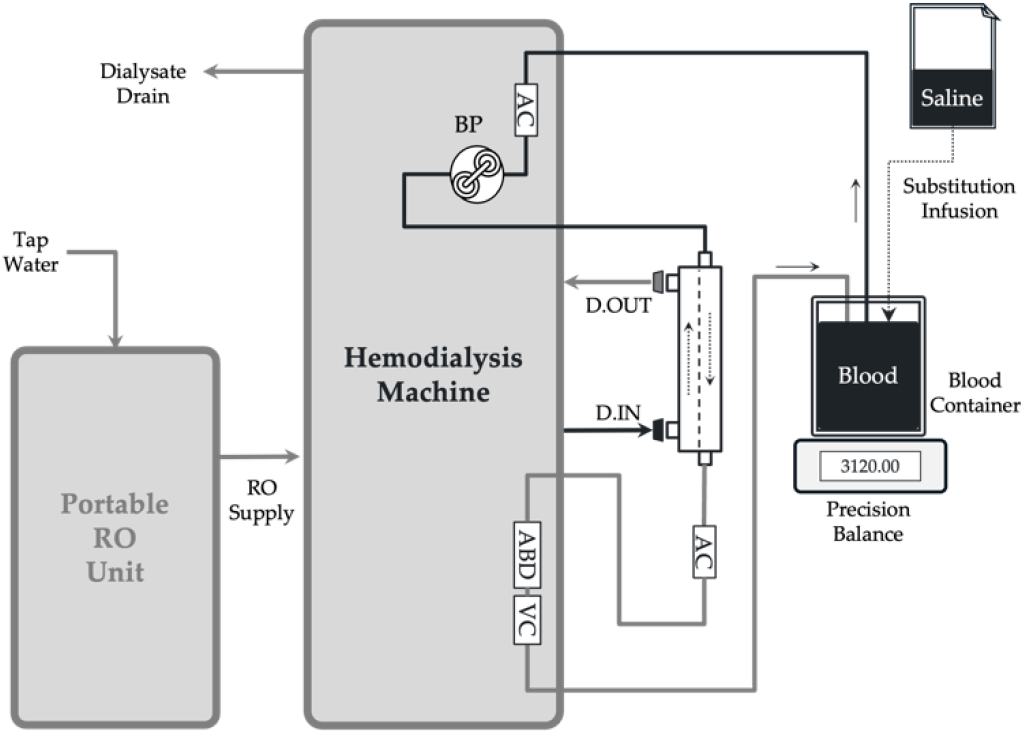
Experimental Circuit Diagram. The circuit includes a hemodialysis device that produces fresh dialysate for the hemodialyzer. About 3 liters of fresh bovine blood was used for the experiment. (D. IN, dialysate inlet; D. OUT, dialysate outlet; RO, reverse osmosis)

About 3 liters of fresh bovine blood, anticoagulated with heparin sodium (Huons, Korea) of concentration 5 units per ml of blood, was used as a blood substitute. The heparin was diluted with normal saline as a supporting agent to lower its viscosity and facilitate circulation. The anticoagulation protocol also involved the infusion of heparin (∼4 units/ml) before the beginning of each dialysis session.

Urea (6.42 g, Samchun, Korea) and creatinine (0.6 g, Sigma, USA) were dissolved in 100 ml of saline solution, which was then mixed with blood. Blood hematocrit levels were adjusted to ∼36% throughout the experiments to compare the hemodialytic efficiencies in the XKIDNEY and conventional groups. Fresh dialysate was prepared by mixing acid and bicarbonate solutions (Boryung, Korea) with purified RO water at a ratio of 1:1.26:32.74. Dialysate conductivity was maintained at ∼14 mS/cm before hemodialyzer.

An Elisio 15H hemodialyzer (Nipro, Japan) with a surface area of 1.5 m2 was used for the experimental dialysis sessions. Blood was circulated at 250 ml/min, and dialysate flow rate was maintained at 500 ml/min for four hours. Blood samples were withdrawn from the blood reservoir every hour before (i.e., t=0) and during dialysis sessions (i.e., t=1, 2, 3, and 4) and analyzed for blood urea nitrogen (BUN), creatinine, plasma proteins, and complete blood count (CBC). Total protein and albumin levels were measured using commercial reagents (Sekisui Chemical, Japan) and a biochemical autoanalyzer (AU480, Olympus, Japan) using the biuret reaction method and a bromocresol green (BCG) technique, respectively. Blood urea nitrogen and creatinine concentrations were also evaluated using a commercial diagnostic kit (Sekisui Chemical, Japan) and a biochemical autoanalyzer (AU480, Olympus, Japan). Urea concentrations were determined by the enzyme-based method of Talke and Schubert, and creatinine concentrations were determined using the Jaffe method.^19^ An auto hematology analyzer (BC-5000 Vet, Mindray, China) was used to quantify the CBC.

Solute reduction ratios were then determined using the following equation:

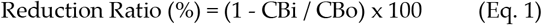

Where, CBo and CBi are solute concentrations at t=0 and after a given number of hours of dialysis, respectively. Solute reduction ratios were used to compare the abilities of XKIDNEY and the conventional device to remove solutes.

### Balancing Error

Prior to beginning each dialysis session, a pseudo-randomized target UF volume (UFt) ranging from 2,400 ml to 3,600 ml was set to the devices, replicating the total 4-hour UF volume typically prescribed by a nephrologist prior to dialysis. Consequently, each experiment featured a distinct target UF volume setting. Then, actual UF volume (UFa) was measured from the blood reservoir using a precision balance during the 4-h HD sessions (FIG. 2). When volume depletion from the blood reservoir due to ultrafiltration reached about 150 ml, an amount of isotonic saline solution equal to the depleted volume was added to the blood reservoir to prevent hemoconcentration. Balancing errors were determined as follows:

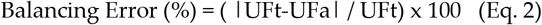

Where UFt is the target UF volume, i.e., the value set for the HD session, and UFa is the actual UF volume measured using the blood container.

### Data Acquisition and Statistical Validation

Arterial blood pressures were measured upstream of the blood pump to ensure adequate blood supply. Venous blood pressures were measured at the venous chamber placed at the same level as the dialyzer blood outlet. Dialysate pressures, temperatures, and conductivities were measured by the XKIDNEY unit. Solutes clearances and reduction ratios during experiments on the XKIDNEY and conventional groups were compared using analysis of variance (ANOVA), and statistical significance was accepted for p values <0.05. Data are expressed as means ± SDs.

## RESULTS

Ex-vivo HD sessions were repeated 19 and 6 times using the XKIDNEY and NCU-18 units, respectively, and no technical malfunctions associated with the XKIDNEY system were observed. Blood and dialysate flow rates were preserved at pre-set values during all HD sessions. In addition, total protein levels, albumin concentrations, and blood hematocrit levels were also preserved throughout dialysis sessions in both groups (Table 1). Solute results revealed that most small uremic molecules were removed from blood within an hour; reduction ratios for urea and creatine were 98.8 to 99.2% and 96.6 to 96.8%, respectively (Table 1).

**TABLE 1.**
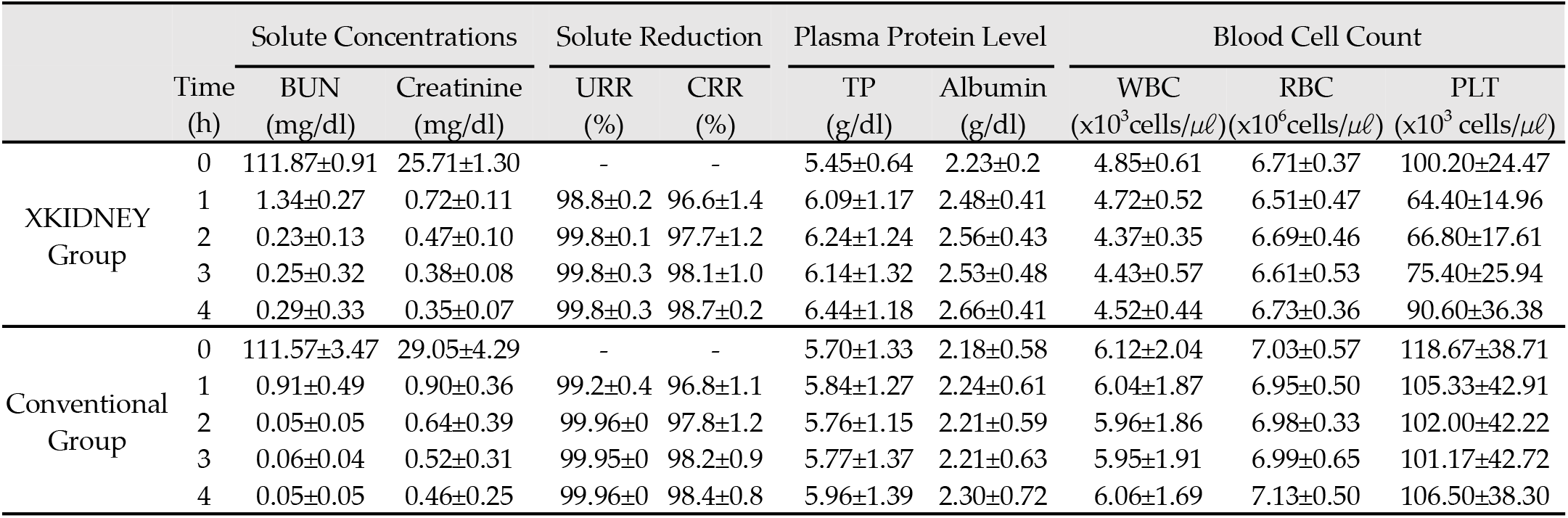
Experimental Results. (BUN, blood urea nitrogen; URR, urea reduction ratio; CRR, creatinine reduction ratio; TP, total protein; WBC, white blood cell; RBC, red blood cell; PLT, platelet)

|  | Time<br>(h) | Solute Concentrations |  | Solute Reduction |  | Plasma Protein Level |  | Blood Cell Count |  |  |
| --- | --- | --- | --- | --- | --- | --- | --- | --- | --- | --- |
| | | BUN<br>(mg/dl) | Creatinine<br>(mg/dl) | URR<br>(%) | CRR<br>(%) | TP<br>(g/dl) | Albumin<br>(g/dl) | WBC<br>($\times 10^3$ cells/$\mu\ell$) | RBC<br>($\times 10^6$ cells/$\mu\ell$) | PLT<br>($\times 10^3$ cells/$\mu\ell$) |
| XKIDNEY<br>Group | 0 | 111.87±0.91 | 25.71±1.30 | - | - | 5.45±0.64 | 2.23±0.2 | 4.85±0.61 | 6.71±0.37 | 100.20±24.47 |
| | 1 | 1.34±0.27 | 0.72±0.11 | 98.8±0.2 | 96.6±1.4 | 6.09±1.17 | 2.48±0.41 | 4.72±0.52 | 6.51±0.47 | 64.40±14.96 |
| | 2 | 0.23±0.13 | 0.47±0.10 | 99.8±0.1 | 97.7±1.2 | 6.24±1.24 | 2.56±0.43 | 4.37±0.35 | 6.69±0.46 | 66.80±17.61 |
| | 3 | 0.25±0.32 | 0.38±0.08 | 99.8±0.3 | 98.1±1.0 | 6.14±1.32 | 2.53±0.48 | 4.43±0.57 | 6.61±0.53 | 75.40±25.94 |
| | 4 | 0.29±0.33 | 0.35±0.07 | 99.8±0.3 | 98.7±0.2 | 6.44±1.18 | 2.66±0.41 | 4.52±0.44 | 6.73±0.36 | 90.60±36.38 |
| Conventional<br>Group | 0 | 111.57±3.47 | 29.05±4.29 | - | - | 5.70±1.33 | 2.18±0.58 | 6.12±2.04 | 7.03±0.57 | 118.67±38.71 |
| | 1 | 0.91±0.49 | 0.90±0.36 | 99.2±0.4 | 96.8±1.1 | 5.84±1.27 | 2.24±0.61 | 6.04±1.87 | 6.95±0.50 | 105.33±42.91 |
| | 2 | 0.05±0.05 | 0.64±0.39 | 99.96±0 | 97.8±1.2 | 5.76±1.15 | 2.21±0.59 | 5.96±1.86 | 6.98±0.33 | 102.00±42.22 |
| | 3 | 0.06±0.04 | 0.52±0.31 | 99.95±0 | 98.2±0.9 | 5.77±1.37 | 2.21±0.63 | 5.95±1.91 | 6.99±0.65 | 101.17±42.72 |
| | 4 | 0.05±0.05 | 0.46±0.25 | 99.96±0 | 98.4±0.8 | 5.96±1.39 | 2.30±0.72 | 6.06±1.69 | 7.13±0.50 | 106.50±38.30 |

Balancing data indicated that both devices exhibited a high degree of balancing accuracy. The UF deviation from the UF target was approximately 12.3 g/h for the XKIDNEY unit, which was substantially lower than the 31.1 g/h UF error observed for the NCU-18 unit. Consequently, the balancing error for the XKIDNEY unit was significantly lower than that of the NCU-18 unit (1.59±1.26% vs. 4.04±1.1%, respectively) (FIG. 3).

**FIG. 3.**
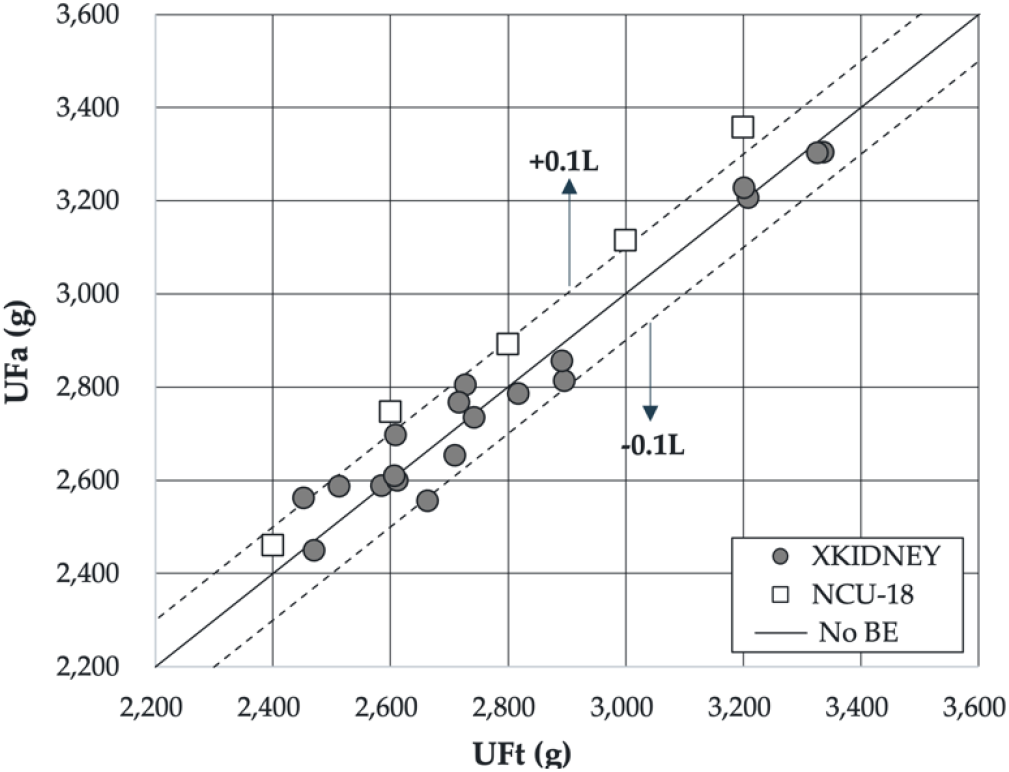
Balancing Data. UFt means the target 4-h ultrafiltration volume, whereas UFa means the actual 4-h ultrafiltration volume. The dotted lines represent volume deviations of +0.1 and -0.1 liters. (BE, balancing error)

## DISCUSSION

Home dialysis provides ESRD patients with physiological and medical benefits. It also provides social benefits, such as more time spent with family members and performing social and professional activities. These medical and social advantages enhance overall quality of life for patients and their families and foster a sense of well-being.^16^

The COVID-19 pandemic disrupted healthcare at an unprecedented level and led to widespread social isolation. The closure of dialysis clinics not only delayed essential HD treatments but forced high-risk patients and their caregivers to visit hospitals frequently, which increased exposure to significant health risks.^20,21^

The advantages of home dialysis are not limited to physiologic and social benefits as they also alleviate the financial burden on the national healthcare systems responsible for providing dialysis. Worldwide, there are ∼3.5 million dialysis patients, and this figure is projected to reach 6 million by 2030.^22^ Such a steep increase in kidney failure cases will place an intolerable burden on healthcare systems already experiencing difficulties meeting the costs of dialysis treatment. Home dialysis presents a potential means of mitigating these costs through the redistribution of healthcare labor. Traditional in-center HD treatment requires the direct involvement of healthcare professionals, whereas home dialysis allows patients and their families to assume some of these responsibilities. This redistribution of labor not only reduces hospital costs and increases operational efficiency but also alleviates the financial burden imposed by in-hospital dialysis on national healthcare systems.^23^

Despite its benefits to patients, patient’s families, hospitals, and governments, the conventional bulky, stand-alone HD machines typically used in hospitals are incompatible with the home dialysis treatment paradigm. These machines weigh nearly 120 kg, are complicated to operate, and require an hour of disinfection before and after each session and an in situ water treatment system, which make them impractical for home use. To address these challenges, various efforts have been made to develop portable-sized units, such as Tablo (Outset Medical, California, USA) and SC+ (Quanta, England) units,^24,25^ which provide more practical solutions for home-based HD. The Tablo unit possesses a HD device and a water purification unit, and thus, does not require connection to a dedicated water treatment facility. Therefore, this unit enables greater flexibility because it is relatively easily moved within a hospital to emergency rooms, operating rooms, and dialysis units.^26^ However, it may not be considered portable as it has a working volume of 270 liters and a weight of 88 kg.

The SC+ unit received FDA approval in January 2021. Like a NxStage’s unit, the SC+ unit employs a cartridge-type tubing set, but unlike NxStage that has a peristaltic system within the cartridge, SC+ adopted a pneumatic pump system.^27^ In addition, both systems share the common feature of not requiring separate disinfection of the dialysate compartment due to the adoption of the cartridge-type dialysate tubing set.

However, cartridge-type disposables increase the cost resulting from its complexity in the production. Patients receiving HD cannot discontinue it until they receive a kidney transplant. The number of HD patients worldwide is effectively doubling every ten years, and as mentioned above, numbers are expected to reach six million by 2030.^28^ Therefore, the healthcare budgets of countries already grappling with the cost of HD will face an additional financial burden in the near future. In this context, cartridge-type disposable sets are more complex and have significantly higher production costs than typical tubing sets. Additionally, disposable cartridge sets consume large amounts of plastic, making them undesirable from an environmental perspective.^29^

The devised XKIDNEY unit features substantial flexibility and is compatible with conventional consumables. Furthermore, despite its innovative portability (size and weight) and suitability for various treatment options, it provides cost benefits, resulting from its unique design eliminating the need for elements considered critical for conventional units, such as the balancing chambers and a separate metering UF pump.

HD constitutes a primary means of removing excess body fluid that has accumulated due to kidney failure. Consequently, accurate fluid management (also termed balancing accuracy) is a critical feature of the design of XKIDNEY because it maintains patient hydration levels, ameliorates potential cardiovascular side effects, and corrects abnormalities in plasma composition. The balancing results obtained in this study emphasize the stability of the XKIDNEY unit, which had an average balancing error of 1.59% of the prescribed UF target, matching or surpassing that of the latest HD device.

Further investigations on the developed XKIDNEY unit are warranted. The ex-vivo tests performed in this study were limited in terms of uremic molecule removal. In clinical practice, uremic solutes are distributed throughout the body, whereas in the present study, uremic molecules were added to a few liters of blood,^30^ and as a result, we observed complete uremic molecule removal. Also, we suggest animal studies be conducted to investigate reductions of medium molecular weight solutes *in vivo*. XKIDNEY is designed to provide push/pull HDF, which may be advantageous for removing medium-to large-sized uremic molecules.

## CONCLUSIONS

The XKIDNEY hemodialysis unit provides portability and versatility due to its innovative design, which markedly reduces unit size and weight but maintains fluid management accuracy. XKIDNEY offers a new means of performing HD hemodialysis at home, providing patients with an improved treatment experience. We aim to provide ESRD patients faced with lifelong dialysis an improved form of treatment and tangible support throughout their long treatment journey.

